# Distinctive *in vitro* phenotypes in iPSC-derived neurons from patients with gain- and loss-of-function *SCN2A* developmental and epileptic encephalopathy

**DOI:** 10.1101/2023.02.14.528217

**Authors:** Miaomiao Mao, Cristiana Mattei, Ben Rollo, Sean Byars, Claire Cuddy, Geza Berecki, Jacqueline Heighway, Svenja Pachernegg, Trevelyan Menheniott, Danielle Apted, Linghan Jia, Kelley Dalby, Alex Nemiroff, Saul Mullen, Christopher A. Reid, Snezana Maljevic, Steven Petrou

## Abstract

*SCN2A* encodes Na_V_1.2, an excitatory neuron voltage-gated sodium channel and major monogenic cause of neurodevelopmental disorders, including developmental and epileptic encephalopathies (DEE) and autism. Clinical presentation and pharmocosensitivity vary with nature of *SCN2A* variant dysfunction with gain-of-function (GoF) cases presenting with pre- or peri-natal seizures and loss-of-function (LoF) patients typically having infantile spasms after 6 months of age. Here, we established and assessed patient induced pluripotent stem cell (iPSC) - derived neuronal models for two recurrent *SCN2A* DEE variants with GoF R1882Q and LoF R853Q associated with early- and late-onset DEE, respectively.

Patient-derived iPSC lines were differentiated using a Neurogenin-2 overexpression yielding populations of cortical-like glutamatergic neurons. Electrophysiological and transcriptomic profiles were assessed after 2-4 weeks in culture. Increased neuronal activity at both cellular and network level was observed for R1882Q iPSC-derived neurons at three weeks of differentiation. In contrast, R853Q neurons showed only subtle changes in excitability after four weeks *in vitro*. In alignment with the reported efficacy in some GoF *SCN2A* patients, phenytoin (sodium channel blocker) reduced excitability of neurons to the control levels in R1882Q neuronal cultures. Transcriptomic alterations in neurons were detected for each variant and convergent pathways pointed at the shared mechanisms underlying *SCN2A* DEE.

## Introduction

Developmental and epileptic encephalopathies (DEE) are a group of rare but severe disorders characterized by pharmacoresistant seizures, behavioural abnormalities, and neurodevelopmental delay. *SCN2A*, encoding the voltage-gated sodium channel Nav1.2, has been increasingly implicated in various forms of DEE (1–3). Distinct phenotypes of DEE and responsiveness to sodium channel blockers (SCBs) correlate with the age of seizure onset (3–5). Here, we have studied two *de novo SCN2A* variants, R1882Q and R853Q, which are among the most recurrent *SCN2A* variants causing DEE with an early- (<3 months) and late- (>3 months) seizure onset, respectively.

The R1882Q variant gives rise to a particularly severe form of DEE. The patients experience pre- or peri-natal seizures (“early-onset”), which occur at high-frequency and are accompanied by movement disorders and intellectual disability (4,5). As observed in other patients with early-onset *SCN2A* DEE, treatment with SCBs could improve seizures, but no improvement was seen in associated comorbidities (4–6). Electrophysiological recordings in heterologous expression systems reveal a GoF phenotype for the R1882Q variant (7–9). A mouse model carrying this variant at the corresponding position showed a severe epileptic phenotype with heterozygous mice developing seizures as early as day 1 after birth and dying within the first month of life (10).*Ex vivo* electrophysiological recordings in cortical pyramidal neurons of heterozygous mice revealed an increase in neuronal firing, supporting the findings in heterologous cell lines (10).

Conversely, patients with the *de novo* missense *SCN2A* variant R853Q present with late-onset (>3 months) seizures with neurodevelopmental delay are resistant to treatment with SCBs or other anti-epileptic drugs (4–6). A LoF has been reported as the principle biophysical mechanism for this variant (7,8,11).

The technology based on induced pluripotent stem cells (iPSCs) has emerged as a unique tool for developing DEE patient-specific models to investigate the disease mechanisms during early development. Patient iPSC-derived models are able to recapitulate disruptive phenotypes at cellular or circuit levels, including alteration in cell morphology (12,13), electrophysiological properties (14–16), synapse formation (14,17,18), and neuronal migration (19). Interestingly, some models have also provided insights into disease mechanisms (20) and subsequently, a partial but significant phenotypic rescue by specific anti-seizure medications (12,18,19). However, while the impact of genetic variation on the protein function has been extensively studied in many genetic epilepsies, the genotype-phenotype relatipnship at the individual level has not been bridged. Accordingly, the mechanisms leading to disturbed neurodevelopment remain elusive, potentially limiting the development of urgently needed effective treatments.

In this study, we aim to establish and analyze patient-derived iPSC models representing two major groups within *SCN2A* DEE to provide a valuable platform to investigate disease mechanisms and test and develop novel therapies.

## Results

### Generation of iPSCs and corresponding isogenic control lines from patients with R1882Q and R853Q SCN2A variants

We generated iPSC lines using peripheral blood mononuclear cells (R1882Q) and skin fibroblasts (R853Q) obtained from patients with the diagnosis of *SCN2A* DEE (5,7). The studied variants replace arginine (R) with glutamine (Q) at positions 1882 in the globular C-terminal Domain, and 853 in the fifth membrane-spanning segment of Domain 4 of Na_v_1.2 (Figure 1A). Sanger sequencing confirmed the presence of heterozygous nucleotide substitution (G>A) at the corresponding positions in the patient iPSC lines (Figure 1B, top). In each patient line, arginine homozygosity was restored following CRISPR-mediated correction for the generation of isogenic control cell lines (Figure 1B, top). Both patient cell lines and corresponding isogenic controls maintained a typical human stem cell morphology during culture (Figure 1B, bottom) and showed the expression of pluripotency markers NANOG and TRA160R (Figure S1).

**Figure 1.**
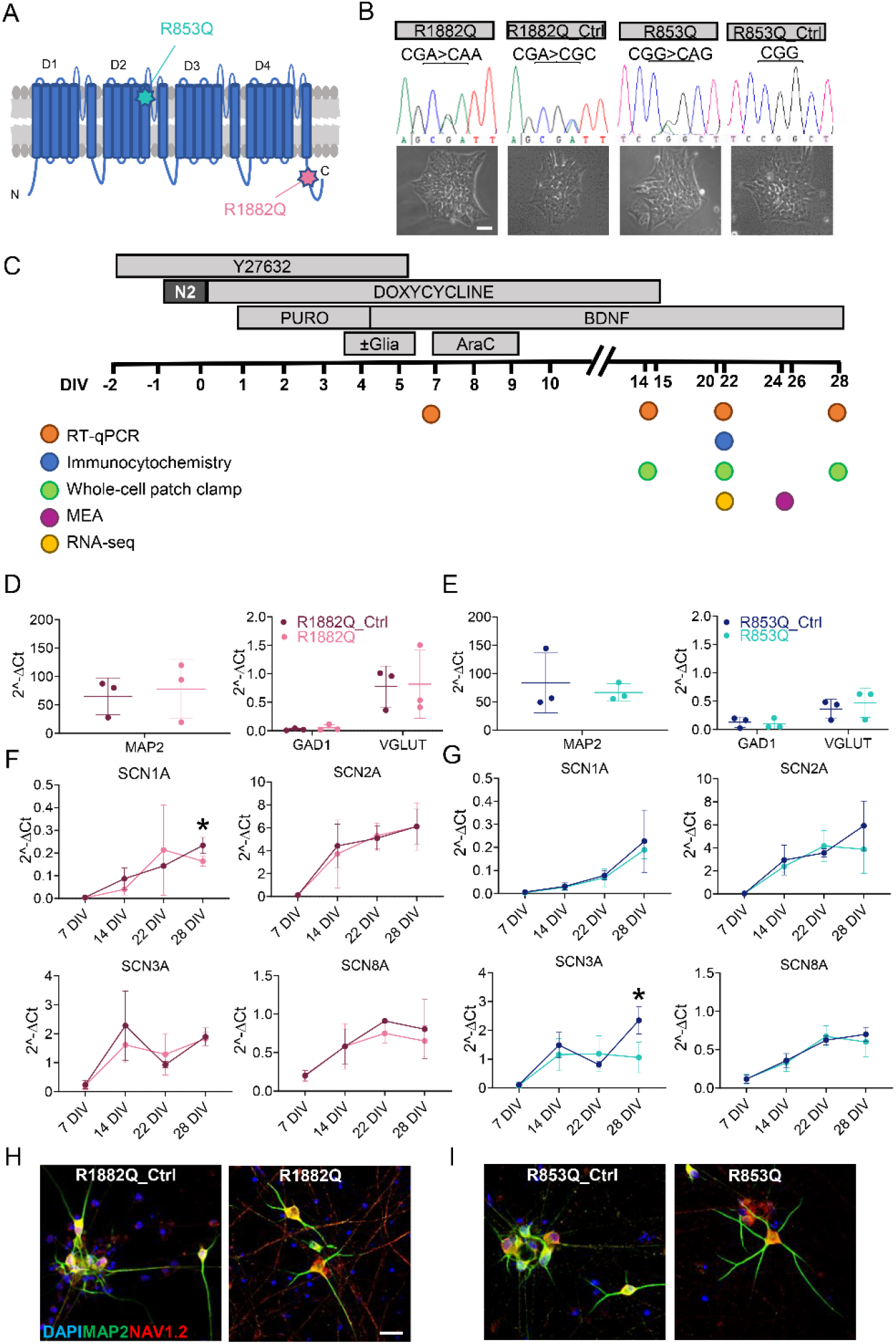
Neuronal differentiation of iPSCs and expression profile analysis of derived cortical-like neurons. (A) Schematic of Na_v_1.2 protein showing the position of R853Q and R1882Q mutations (D, Domain; N, N-Terminal; C, C-Terminal). (B) Sanger sequencing of R1882Q and R853Q iPS cell lines and corresponding CRISPR-mediated corrected cell lines (top) maintained normal morphology (bottom). (C) Schematic overview of the *NGN2* differentiation protocol with details of crucial molecules exposure and analysis timeline. RT-qPCR analysis of neuronal maturation marker MAP2 and cell identity markers GAD1 and VGLUT at 22 DIV in R1882Q (D) and R853Q (E) iPSC-derived neurons and corresponding control. RT-qPCR analysis of sodium channel genes *SCN1A, SCN2A, SCN3A, SCN8A* in R1882Q (F) and R853Q (G) iPSC-derived neurons and corresponding control. Data presented as mean ± SD (n=3 experimental biological replicates with n=3 samples per experiment). Unpaired t-test was used to assess statistical significance. Immunofluorescence analyses of MAP2 and SCN2A expression in R1882Q (H) and R853Q (I) iPSC-derived neurons and corresponding control at 22 DIV. Scale bars, (B) 50 μm, (F, G) 20 μm. \**p*<0.05.

### Derivation of cortical excitatory neurons from patient and control iPSC lines

Because *SCN2A* is robustly expressed in cortical excitatory neurons, which are presumably most affected in *SCN2A* DEE (21), we employed the lentiviral-based *NGN2* differentiation method whereby the upregulation of *NGN2* in iPSCs induces a rapid and reproducible fate toward cortical like-neurons (22). Protocol overview and time points for different phenotypic readouts are presented in Figure 1C. Using RT-qPCR analysis, we showed that neurons obtained from both patient and control iPSCs showed a notable expression of neuronal maturation marker MAP2 after 21 days *in vitro* (DIV). Further analysis to the presence of specific cell identity markers revealed that *NGN2*-derived neurons show a moderate expression of excitatory marker vGLUT1 as well as a detectable expression of inhibitory marker GAD1 without any significant difference between patient iPSC-derived neurons and their corresponding isogenic controls (Figure 1D, E). Taken together, these results confirmed that the cortical identity and maturation of neuronal cultures was unaffected by the genotype.

Subsequently, we interrogated the expression of *SCN2A* and the other brain voltage-gated sodium channel genes *SCN1A, SCN3A* and *SCN8A*. This was done to both validate the *NGN2* neurons as a suitable model for *SCN2A* disorders as well as to evaluate if potential differences in the expression of other sodium channel genes may be contributing to the *SCN2A* disease phenotype. RT-qPCR analysis showed that at early differentiation time points, cells already express all *SCNxA* genes studied with an increasing trend over time up to 28 DIV suggesting that *NGN2*-derived neurons might display time-dependent electrophysiological competence and action potential firing (Figure 1F, G). *SCN1A* expression was significantly higher in the isogenic control compared to the R1882Q patient line at 28 DIV, while the mRNA expression of the other genes did not reach any significant difference across all time points (Figure 1F). Similarly, for the R853Q variant, three of the four genes showed no difference in the expression profile between the patient and the isogenic control line at any of the time points. However, *SCN3A* had a two-fold higher expression in the isogenic control derived neurons at 28 DIV (Figure 1G). Overall, results of our RT-qPCR analysis suggested that in our model, *SCN2A* R1882Q and R853Q variants do not correlate with any transcriptional alteration of *SCN2A* expression and that only *SCN1A* for the R1882Q and *SCN3A* for R853Q variant had higher expression in corresponding controls at four weeks of differentiation. Interestingly, when we interrogated the expression of a specific adult variant of *SCN8A*, the 18A transcript, which is known to encode for the fully functional protein in the adult brain(23,24), we could not detect any expression (data not shown, Ct undetectable). This indicated that *NGN2*-derived neurons, both patient and corresponding control cell lines, only express the *SCN8A* 18N transcript which encodes for the neonatal truncated protein, indicating the foetal nature of our modelling system in the time frame studied. Using immunocytochemistry, we were able to detect the expression of Na_v_1.2 protein in neurons at DIV 22 (Figure 1H, I).

These findings represent a validation of the *NGN2*-derived neurons for modelling *SCN2A* variants and present the basis for the subsequent electrophysiological and transcriptomic analysis.

### R1882Q patient iPSC-derived neurons have a clear hyperexcitable phenotype

Since the expression of brain sodium channels at the mRNA level appeared to have stabilised after three weeks in culture (Figure 1F, G), and the differentiated neurons presented with consistent firing at that age, we performed whole-cell patch clamp recordings at 20-22 DIV (Figure 2; Table 1). No significant differences in passive membrane properties, including cell capacitance, input resistance and resting membrane potential, or in spontaneous firing rates was found between the R1882Q and isogenic control neurons (Figure 2A). However, action potential (AP) firing was increased in patient cells over a range of current injections (Figure 2B-C), with a corresponding significant increase in maximum AP frequency (Figure 2D, left) but no difference in rheobase (Figure 2D, right). Only 3/61 control neurons and 0/49 R1882Q neurons firing <10 AP per second (Figure 2E) at this time point. We assessed single AP properties and found significant differences in AP threshold, peak amplitude, and AP rise time, with no change in AP decay time (Figure 2F), consistent with a sodium channel GoF leading to hyperexcitable neuronal phenotype in the patient cell line. For a subset of electrophysiological experiments, we generated co-cultures of *NGN2* neurons and human astrocytes, which were analyzed using multielectrode array (MEA) recordings (Figure S2). Patient iPSC-derived neurons showed increased mean firing and network burst rates (Figure S2C, D). Analysis of parameters used to assess the synchronicity of neuronal activity, such as jitter or kappa, showed no significant differences (Figure S2E, F). Taken together, these electrophysiological changes indicate an increased neuronal excitability in neural cultures derived from patients with the *SCN2A* R1882Q variant.

**Figure 2.**
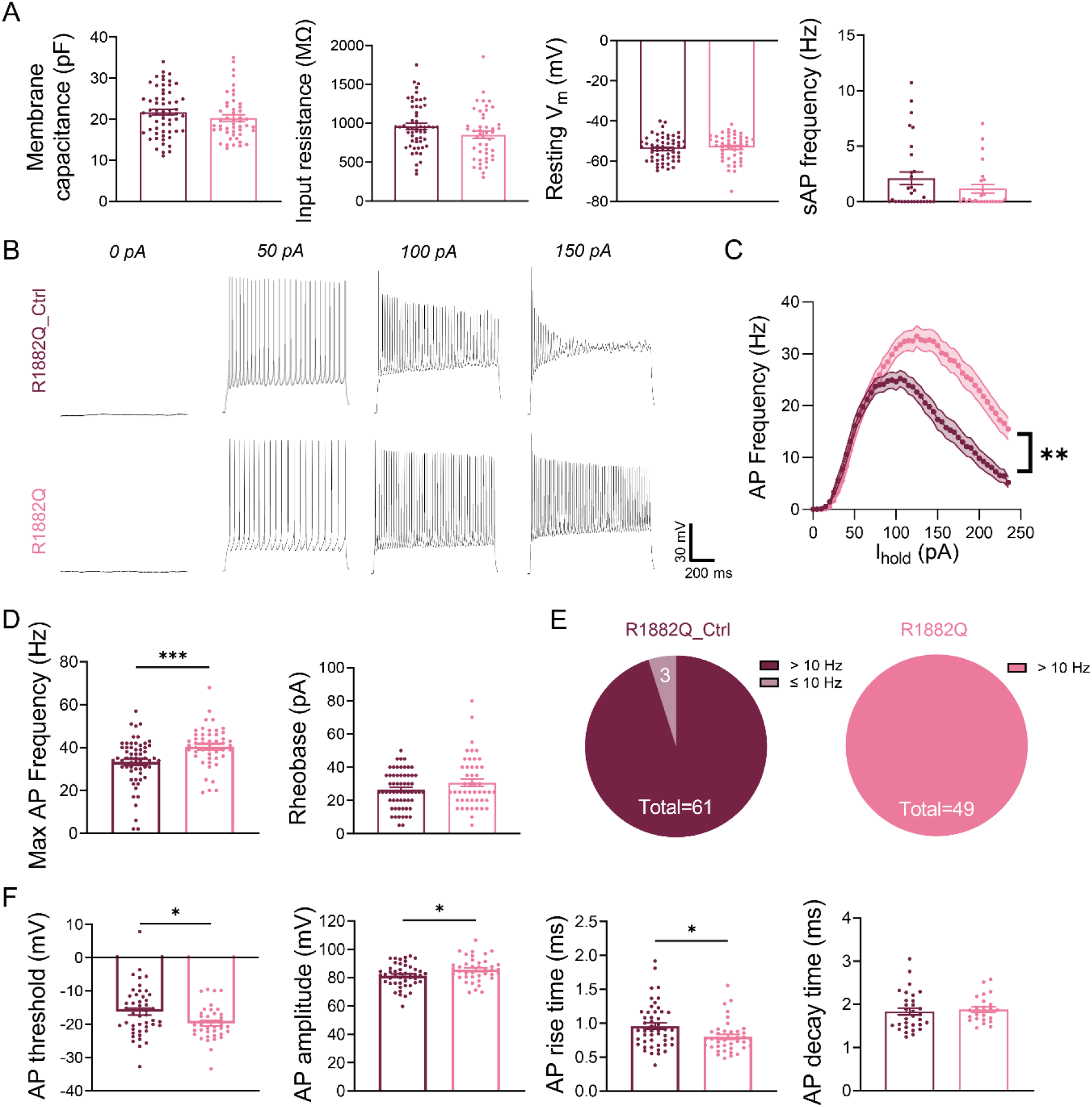
Whole-cell patch clamp recordings of R1882Q patient and isogenic control cell lines at DIV 20-22. (A) Passive membrane properties (from left to right): membrane capacitance, input resistance, resting membrane potential and spontaneous AP frequency. (B) Representative traces of current clamp recordings. (C) Mean input-output function of AP firing in response to current injection (R1882Q isogenic control, n=61; R1882Q, n=48), *p*<0.01 (minimum *p* value, two-way ANOVA with multiple comparisons test). (D) Maximum AP frequency (left, *p*=0.0005) and rheobase (right). (E) Pie charts showing proportion of cells with >10 Hz or ≤10 Hz maximum AP frequencies. (D-F) AP properties analyzed for the second AP at 5 pA above rheobase. (F) Single AP properties (from left to right): AP threshold (*p*=0.0155), AP peak amplitude (*p*=0.0245), AP 10-90% rise time (*p*=0.0108) and AP 90-10% decay time. Unpaired Mann-Whitney tests (A, E and F). Mean±SEM.

### R853Q patient iPSC-derived neurons show a subtle electrophysiological phenotype

In contrast to R1882Q, R853Q patient cells did not show any clear differences in AP firing or single AP properties relative to the isogenic control at DIV 20-22 (Figure 3 and Table 1), except a small increase in the proportion of low-firing cells in the R853Q cell line (Figure 3D). We additionally assayed the R853Q patient and control cells at two other time points (Figure 4). While no differences in firing rates were seen at DIV 13-15 (Figure 4A), there were more low-firing cells present in the R853Q cell line than in control (Figure 4B). In addition, R853Q patient cells had larger membrane capacitance potentially indicating larger membrane surface (Figure 4C). We performed a morphological assessment revealing no significant differences in the soma size between the patient and control cells at this time point (Figure 4D). However, we could observe greater proportions of R853Q patient cells with medium- (81-111 μm^2^) and large- (112-166 μm^2^) sized soma compared to R853Q isogenic control cells (43.9% vs 34.1% for medium-sized soma, 17.6% vs 13.7% for large-sized soma, respectively). At a later time point (DIV 28-31), R853Q patient cells had similar firing evoked by current injections compared to the control (Figure 4E) and similar number of low-firing cells (Figure 4F). Interestingly, R853Q cells displayed more depolarised resting membrane potential (Figure 4G) and fired more spontaneous APs (Figure 4H).

**Figure 3.**
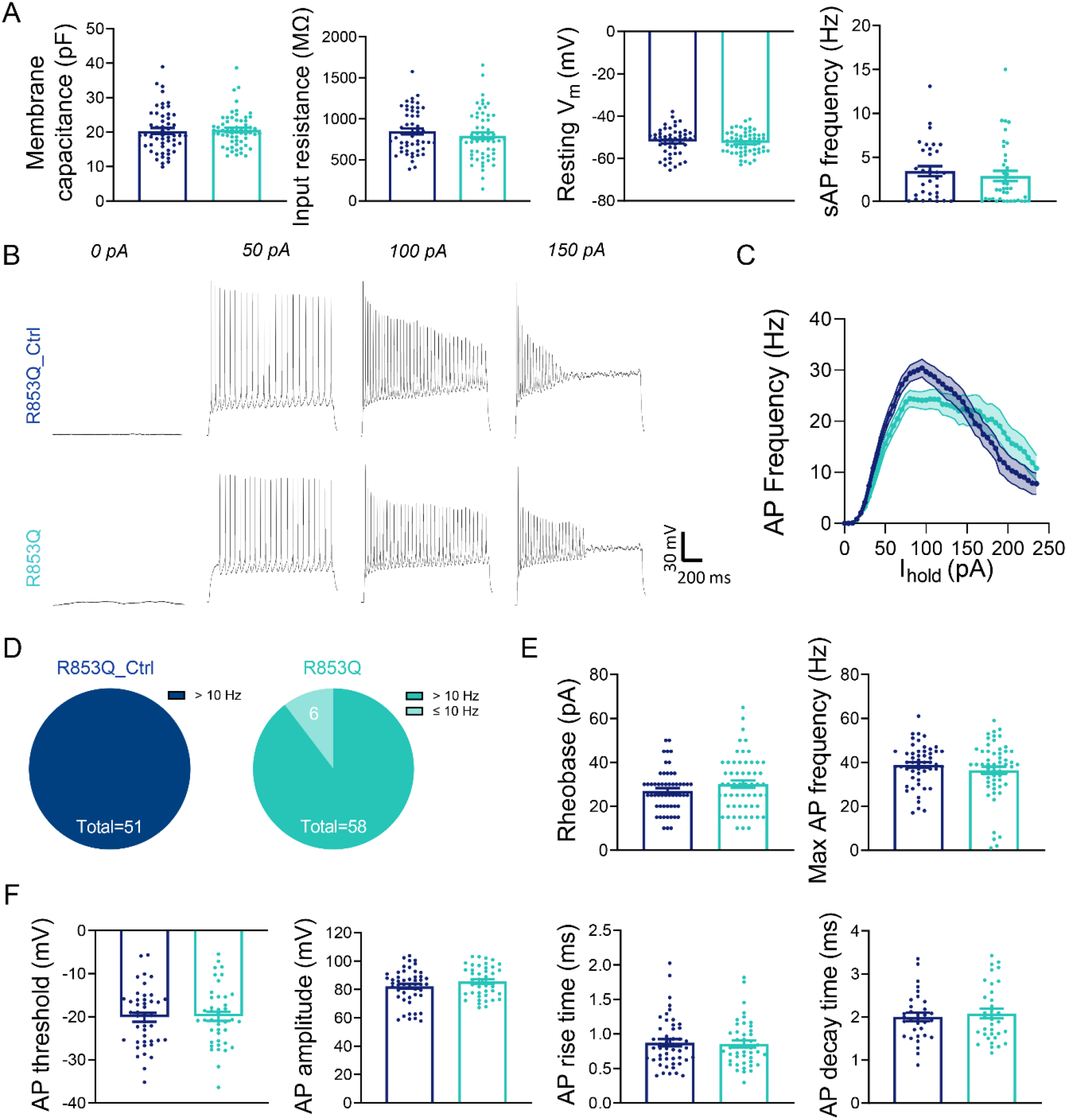
Whole-cell patch clamp recordings of R853Q patient and isogenic control cell lines at DIV 20-22. (A) Passive membrane properties (from left to right): membrane capacitance, input resistance, resting membrane potential and spontaneous AP frequency. (B) Representative traces of current clamp recordings. (C) Mean input-output function of AP firing in response to current injection (R853Q isogenic control, n=54; R853Q, n=58). (D) Pie charts showing proportion of cells with >10 Hz or ≤10 Hz maximum AP frequencies. (E) AP firing properties (from left to right): rheobase and maximum AP frequency. (F) Single AP properties (from left to right): AP threshold, AP peak amplitude, AP 10-90% rise time and AP 90-10% decay time. Unpaired Mann-Whitney tests (A, E and F). Mean±SEM.

**Figure 4.**
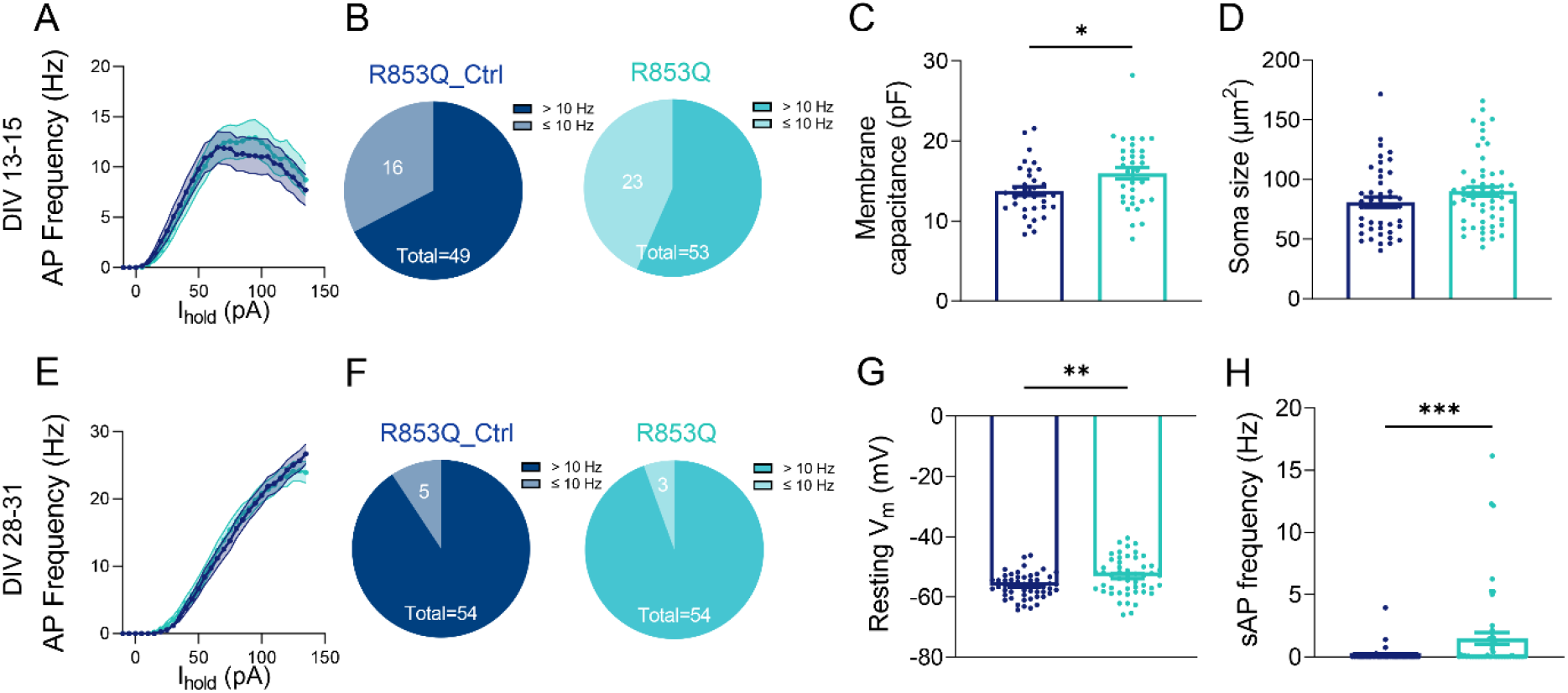
Whole-cell patch clamp recordings of R853Q patient and isogenic control cell lines at DIV 13-15 and DIV 28-31. (A-D) DIV 13-15. (E-H) DIV 28-31. (A, E,) Mean input-output function of AP firing in response to current injection at DIV 13-15 (A) and DIV 28-31 (E). (B, F) Pie charts showing proportion of cells with >10 Hz or ≤10 Hz maximum AP frequencies at DIV 13-15 (B) and DIV 28-31 (F). (C) Membrane capacitance at DIV 13-15 (*p*=0.026). (D) Soma size at DIV 13-15. (G) Resting membrane potential at DIV 28-31 (*p*=0.0032). (H) Spontaneous AP frequency at DIV 28-31 (*p*=0.0002). Unpaired Mann-Whitney tests (C, D, G and F). Mean±SEM.

Analysing the *in vitro* development of these cell lines across all three time points (Figure S3) revealed a difference in the trajectory of resting membrane potential (Figure S3B) between R853Q and its isogenic control, while no significant differences were seen in the developmental trajectory of AP firing or single AP properties over the course of 2 weeks (Figure S3).

### Phenytoin modulates AP firing in R1882Q and R853Q patient cells

Phenytoin, a sodium channel blocker, has been widely used in the treatment of epileptic seizures (4,5,25). It has been well-reported that DEE patients with gain- or LoF *SCN2A* variants respond differentially to phenytoin treatment (4,5,7,26). Therefore, we set out to test the effects of phenytoin on neuronal excitability in the *SCN2A* DEE patient iPSC-derived neurons (Figure 4). As expected, phenytoin caused a decrease in AP firing in both patient cell lines in a dose-dependent manner. Notably, 30 μM phenytoin could effectively reduce AP firing in R1882Q patient cells to a level similar to the untreated isogenic control (Figure 4 B-D). In contrast, phenytoin treatment in neurons derived from R853Q patient decreased AP firing further away from the untreated isogenic control (Figure 4 E-G). These observations are consistent with the reported clinical outcomes of phenytoin treatment in *SCN2A* DEE patients, with GoF *SCN2A* mutations, while being ineffective or even worsening the conditions in those with LoF mutations (4,5,7,26).

### Transcriptomic signatures of SCN2A DEE

To identify genes and molecular pathways dysregulated in *SCN2A* early- and late-onset DEEs, we then performed bulk RNA sequencing of R1882Q and R853Q patient-derived and isogenic controls neurons at DIV 21. Principal component analysis (PCA) indicated that the R1882Q variant was responsible for more variance in the generated data set than R853Q variant, with tight clustering between biological replicates in patient compared to control groups (Fig S5 and Table 3). Accordingly, differential gene expression analysis identified 406 and 83 significantly up- and down-regulated genes for R1882Q- and R853Q-derived neurons compared to their isogenic controls, respectively, at 5 % false-discovery rate (FDR) (Figure 5 A, B). Interestingly, the majority of genes (356 of 406 genes) were upregulated in the R1882Q neurons, whereas R853Q neurons had a similar percentage of up- and down-regulated genes (40 of 83 genes up-regulated, and 43 down-regulated).

**Figure 5.**
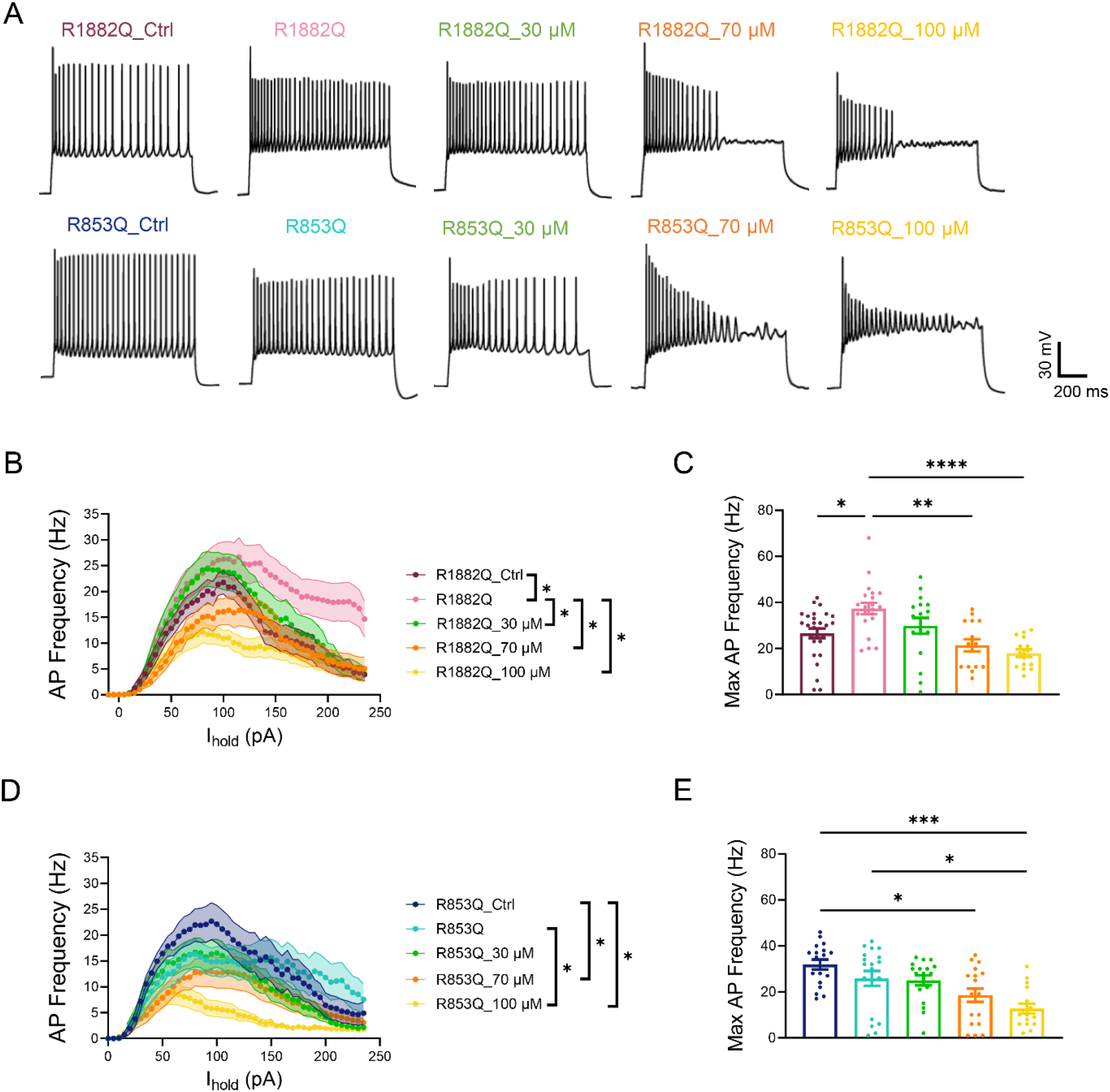
Phenytoin reduces AP firing in both R853Q R1882Q patient iPSC-derived neurons in a dose-dependent manner. (A) Representative traces of current clamp recordings with 100 pA current injections. (B-C) Summary data of AP firing in R1882Q neurons. (B) Mean input-output function of AP firing in response to current injection (R1882Q isogenic control vehicle, n=27; R1882Q vehicle, n=21; R1882Q 30 μM phenytoin, n=17; R1882Q 70 μM phenytoin, n=15; R1882Q 100 μM phenytoin, n=15), *p*<0.05 (two-way ANOVA with multiple comparisons test). (C) Maximum AP frequency (one-way ANOVA with Tukey’s multiple comparisons test, \**p*<0.05, \*\**p*<0.01 and \*\*\*\**p*<0.0001). (D-E) Summary data of AP firing in R853Q neurons. (D) Mean input-output function of AP firing in response to current injection (R853Q isogenic control vehicle, n=18; R853Q vehicle, n=19; R853Q 30 μM phenytoin, n=19; R853Q 70 μM phenytoin, n=18; R853Q 100 μM phenytoin, n=16), *p*<0.05 (two-way ANOVA with multiple comparisons test). (E) Maximum AP frequency (one-way ANOVA with Tukey’s multiple comparisons test, \**p*<0.05, \*\*\**p*<0.001). Error bars represent SEM.

**Figure 6:**
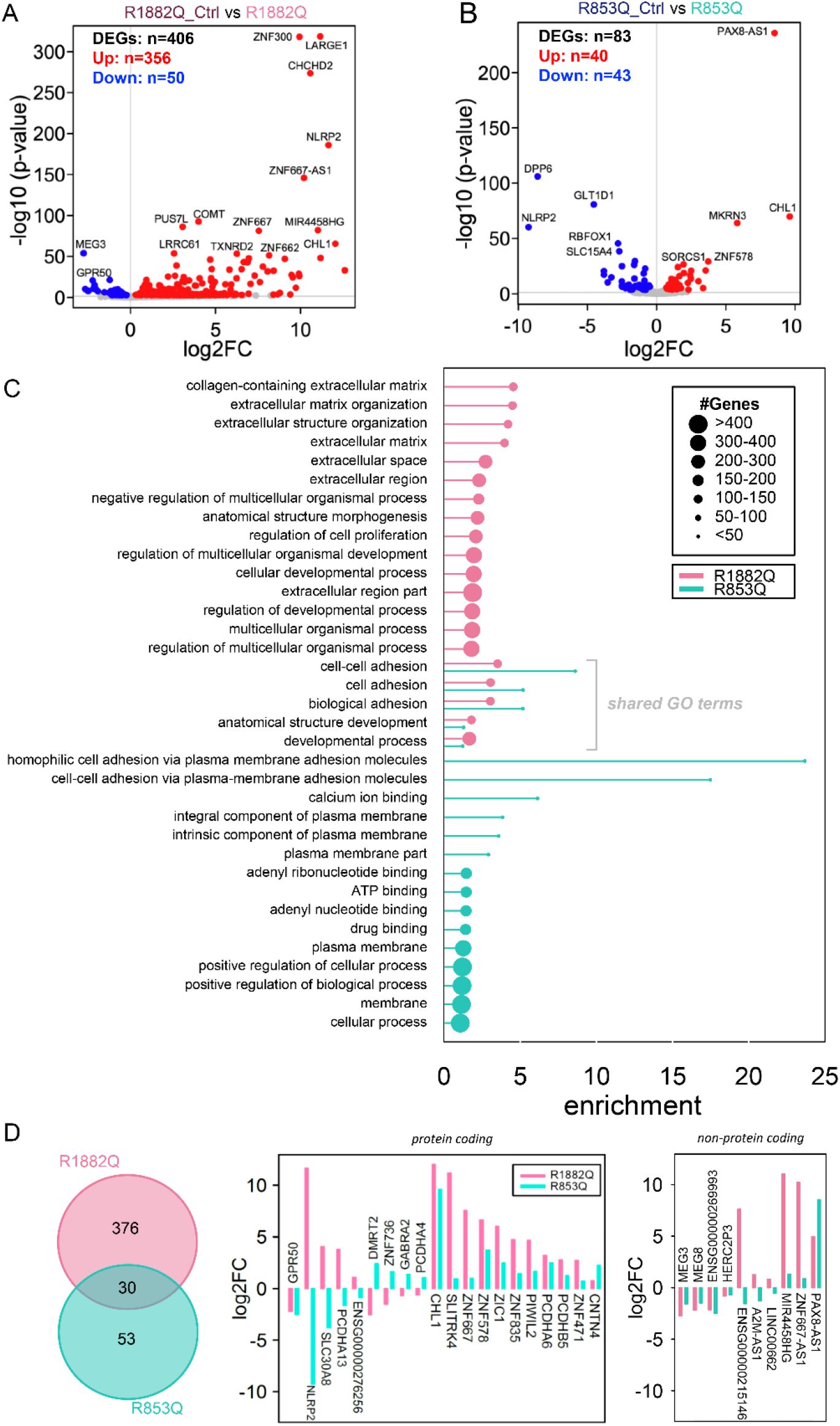
mRNA-sequencing of *SCN2A* R1882Q and R853Q derived *NGN2* neurons revealed unique and convergent DEGs and biological pathways. Volcano plots of significantly differentially expressed (FDR q<0.05) genes (DEGs) in (A) *NGN2* R1882Q and (B) R853Q-derived neurons versus corresponding isogenic controls. Significantly up- and down-regulated genes are shown in red and blue respectively. (C) Lollipop plot showing the top 20 significantly enriched (FDR q<0.05) gene ontology (GO) terms for both variants. Overlapping terms are grouped in the centre. The enrichment values are based on the standard hypergeometric distribution approach described in (63), with the circle size represented by *n*, the number of DEGs that were highly ranked (by small FDR q-value) and represented in each GO term. (D) 30 overlapping DEGs are identified between the two variants. Bar plots of *log_2_FC* values indicate which were up- or down-regulated in each variant as well as whether they were (non)protein-coding.

Gene ontology (GO) analysis of biological processes indicated that *SCN2A* R1882Q variant alters the expression of genes involved in regulation of multicellular organismal processes, developmental processes, and extracellular structure organization, while *SCN2A* R853Q variant alters the expression of genes involved in regulation of biological processes, plasma membrane, ATP and drug binding (Figure 5C). Although many different GO functions were unique to individual datasets, there were also some related functions among the datasets, including cell adhesion and development processes. These fundings prompted us to further assess convergent gene pathways between early- and late-onset DEE *SCN2A* variants.

### Biological convergence of early and late SCN2A DEE

Comparison of differentially expressed genes (DEGs) in R1882Q and R853Q revealed 30 overlapping dysregulated genes (Figure 5D), with 10 of these being non-protein coding. Many of the 20 protein-coding overlapping DEGs (12 of 20 genes) were dysregulated in same direction in both data sets. The predominant effect in this group was up-regulation (11 out of 12), with the largest increase seen for *CHL1*. The only downregulated gene was *GPR50*. Among the shared protein-coding DEGs showing opposite dis-regulation (8 of 20 genes), half were up-regulated in R1882Q and down-regulated in R853Q and the other half were changed in the opposite direction. Within this gene pool, *NLRP2* expression displayed the most severe dysregulation of a similar degree but in the opposite direction between the two variants.

## Discussion

In this study, we have generated iPSC lines from two patients carrying some of the most recurrent *SCN2A* variants causing early (R1882Q) or late (R853Q) seizure onset DEE. A CRISPR/Cas9 approach was used to generate corresponding isogenic controls. We employed a quick neuronal differentiation protocol (*NGN2* overexpression) to produce cortical-like excitatory neurons which were then assessed within 4 weeks of differentiation using patch clamp analysis, MEA, RT-qPCR and RNA sequencing. We demonstrate that the R1882Q patient iPSC-derived neurons had increased excitability, consistent with the previously reported GoF phenotype seen in heterologous expression systems and *in silico* predictions (7–9). In contrast, patient iPSC-derived neurons modelling LoF R853Q variant had a more subtle electrophysiological phenotype, dependent on the time point of the analysis. In addition to functional assessment, RNA sequencing identified higher numbers of differentially expressed genes (DEGs) for the R1882Q line vs isogenic control compared to R853Q. Interestingly, some of the alterations in the gene expression affected a pool of identical genes pointing that shared mechanisms could underlie two distinctive forms of *SCN2A* DEE.

Sodium channels are critical for the functionality of generated neurons. We assessed the expression of the major brain sodium channels in the two isogenic pairs and could detect no significant changes in expression at the time points studied, except for a mildly decreased expression of *SCN1A* in R1882Q neurons and *SCN3A* in R853Q neurons at latest time point studied. While at this stage the expression of *SCN1A* in the *NGN2* neurons is still very low, *SCN3A* has been suggested as the predominant sodium channel (27,28), indicating that its reduced expression in R853Q neurons could be a relevant molecular mechanism. In addition to its canonical role in regulating sodium conductance in cortical progenitors during neurogenesis, *SCN3A* has been indicated as a possible candidate in regulation of neuronal development through action potential-independent mechanisms. In fact, altered expression of *SCN3A* has been associated with disruption of neurite branching, neural migration and cerebral cortex folding (29,30). Accordingly, reduced *SCN3A* expression in R853Q neurons could impact development and maturation of neurons independent of their sodium conductance and thus lead to DEE.

Seven patients carrying *de novo* R1882Q mutation in *SCN2A* causing early-onset (0-3 months) DEE have been identified to-date (7). Using patch clamp and MEA recordings we showed an increase in neuronal excitability in patient iPSC-derived neurons compared to isogenic controls at DIV 21. Recently, a different *SCN2A* early-onset DEE variant (L1342P) was shown to cause channel GoF in mammalian cell expression system and hyperexcitability in human iPSC-derived neuronal cultures using whole-cell patch clamping and MEA recordings (31). The mutation was genetically engineered into a donor/control cell line and characterized using a dual-SMAD neuronal differentiation protocol. In contrast, we used patient-derived iPSC line and an isogenic control line in which the variant is “corrected”, enabling the assessment of its functional effects and personalised drug responsiveness in the patient-specific genetic background. In addition, the use of *NGN2* differentiation protocol provides a method to fast-track functional assessments in a more homogeneous neuronal population. Despite the different approaches employed, both studies demonstrate that the GoF *SCN2A* variants lead to increased excitability in human stem cell-derived neuronal models.

Conversely, R853Q patient iPSC-derived neurons showed no significant differences in electrophysiological properties at DIV 21, despite a LoF phenotype described in heterologous expression systems and *in silico* prediction of reduced neuronal firing (7,8,11). However, examination at an earlier (DIV 13-15) and later (DIV 28-31) time points revealed an increase in membrane capacitance and neuronal excitability at rest, respectively. The increase in membrane capacitance at the earlier time point in the R853Q neurons could indicate an abnormal development of cortical excitatory neurons, potentially leading to malformation of neuronal circuitry and subsequently altered brain development. The cell membrane assay did not show a significant increase of soma size in patient cells, but the change in capacitance could also be attributed to changes in the size of neuronal processes, which were not assessed in our study. A more depolarised resting membrane potential at DIV 28-31 suggests reduced maturity of patient iPSC-derived neurons. Furthermore, a more depolarised membrane potential is closer to the threshold of action potential firing, which can increase the spontaneous activity seen in these neurons. Therefore, the patient R853Q iPSC-derived neuronal model suggests that LoF *SCN2A* variant does not simply reduce neuronal excitability but the mechanisms could implicate development and maturation of impacted neurons.

Many of the identified *SCN2A* LoF variants cause developmental delay, intellectual disability and/or mental disorders such as autism and schizophrenia, with or without epileptic seizures (3,5,32–35). Studies in mice with heterozygous deletion of *SCN2A* showed that they partially recapitulate patient phenotypes, including absence seizures, mental disability and hyperactivity (36,37). In addition, Na_v_1.2 has been shown to be important in regulating dendritic excitability and back propagation of APs, critical for synaptic function in a mouse model of *Scn2a* haploinsufficiency (38). Furthermore, a loss of Na_v_1.2 channel function in mouse models can cause neuronal hyperexcitability due to dysregulation of potassium channels and a failure to adequately repolarise neurons during AP firing (39). More complex patient-derived neuronal models may be able to recapitulate some of these findings.

We further examined the effect of phenytoin on neuronal excitability. Phenytoin, as well as other SCBs, have been reported to reduce seizure frequencies in patients with GoF *SCN2A and SCN8A* variants causing severe early-onset seizures (40–46). On the other hand, SCBs are usually not effective in patients with LoF *SCN2A* variants (5,7). In this study, phenytoin (30 μM) reduced high-frequency AP firing caused by the GoF *SCN2A* variant (R1882Q) to the level of the isogenic control. In comparison, the reduction of AP firing by phenytoin in the LoF cell line (R853Q) only increased the difference to isogenic control, which is unlikely to be beneficial.

RNA sequencing revealed important molecular signatures of early- and late-onset *SCN2A* DEEs. Many DEGs were found in R1882Q patient neurons, most of them showing upregulation. While this effect may be due to a robust change in electrophysiological activity of these neurons, we cannot exclude that the DEGs contributed to hyperexcitability. In contrast, fewer DEGs found in R853Q derived neurons, suggest a need for a more sophisticated modelling system. Looking at the gene ontology (GO) terms, the R1882Q variant mostly affected regulation of developmental processes and extracellular structure, whereas R853Q variant altered the regulation of biological processes, plasma membrane, ATP and drug binding. This is an interesting addition to the functional analysis, suggesting that R1882Q increased excitability can be linked to changes in the developmental regulation. In turn R853Q phenotype may reflect more complex metabolic changes affecting neuronal development.

We further asked if *SCN2A* dysfunction, irrespective of gain- or loss-of-function, could affect convergent pathways. Among the GO terms, regulation of cell adhesion which was already described in autism spectrum disorders (47), is reported in both patient lines. Interestingly, 30 overlapping DEGs were found between the two *SCN2A* isogenic pairs, of which 20 were protein-expressing genes, with *NLRP2* and *CHL1* expression showing the most pronounced change. *NLRP2* is altered in opposite directions in the two models (increased expression in R1882Q and decreased in R853Q), whereas *CHL1* is upregulated in both. Dysregulation of *NLRP2*, involved in the regulation of immune responses, has been observed in familial early-onset Alzheimer’s (48), late-onset Alzheimer’s (49,50), and in bipolar disorder (51,52). Deletions of *CHL1* have been described in patients with neurodevelopmental delay and seizures, but less is known about the potential effect of *CHL1* overexpression. Two reported cases with a *CHL1* copy number variation (CNV) presented some overlapping phenotypes including intellectual disability (53,54). Contribution of 10 non-protein-coding genes corresponds to previous studies showing disruptions in non-coding RNAs in ASD subjects (55).

Similarly, to others, we encountered limitations in analysing electrophysiological properties of iPSCs-derived neurons due to the intrinsic variability in maturation within culture as well as across separate biological experiments. Additionally, we appreciate the benefits of using *NGN2* neurons in co-culture with astrocytes (56), which would promote the maturation of *NGN2* neurons and the formation of functional synapses which are not present in pure *NGN2* neuronal cultures (22,57,58). In this study co-cultures were only used for MEA recordings to avoid additional variance introduced to patch clamp recordings. Other *in vitro* modelling approaches, such as cortical organoids, would provide more sophisticated insights in disease mechanisms.

Furthermore, fluorescent reporters expressed under specific promoters to identify neuronal subtypes or functional maturity of neurons would reduce data variance and improve interpretation in patch clamp recordings from 2D or 3D cultures [see (59)].

In conclusion, our findings demonstrate the capability of patient iPSC-derived models to recapitulate crucial aspects of the disease phenotype and pharmacosensitivity for gain and loss of function *SCN2A* variants. The models generated in this study provide novel insights into the disease mechanisms and can aid the development of *in vitro* platforms for identifying subtypes of DEE and development of novel therapies for these patients.

## Methods

### Generation and culture of SCN2A R1882Q and R853Q iPSC lines

This study was approved by the Austin Health Ethics committee, ethics number HREC/16/Austin/472.

The two stem cell lines were generated from peripheral blood mononuclear cells (PBMCs) isolated from cord blood obtained at birth for the *SCN2A* R1882Q individual, and from skin fibroblasts for the *SCN2A* R853Q individual. These PBMC’s and skin fibroblasts were reprogrammed by forced-expression of KSOM factors (Kl4, Sox2, Oct4, C-Myc) introduced by Sendai virus according to the manufacturer’s instructions (Cytotune iPS 2.0 Sendai reprogramming kit, ThermoFisher) or by using an episomal method, respectively (60).

Following isolation of individual colonies from feeder layers, iPS cells were expanded on 5μg/ml vitronectin (#A14700, ThermoFisher), grown in E8 essential medium (#A1517001, E8 medium, ThermoFisher) and passaged using 0.5 mM EDTA (#15575-038, Life Technologies). These cells were referred to as R1882Q and R853Q iPS cells.

### Generation and culture of isogenic corrected control iPS cell lines

*SCN2A* R1882Q and R853Q iPS cells were used as targets for CRISPR mediated correction of corresponding mutation. R853Q correction has already been described (60). For R1882Q, a DNA repair template was constructed comprising approximately 600bp homology arms flanking the patient-specific mutation (NM_021007.2:c.5645G>A) which was cloned into the pSMART-HCKan (#AF532107, GenBank) plasmid vector. A synonymous 3 bp change was incorporated to facilitate the identification of targeted clones and two additional synonymous changes were included to act as Cas9-blocking mutations (Supplementary methods). For targeting, 10×10^6^ iPS cells were harvested with TrypLE (#12604013, ThermoFisher) and electroporated (1100V, 30ms, 1 pulse) using the Neon transfection system (ThermoFisher) and plated over 4 wells of a Matrigel-coated (BD Biosciences) 6-multiwell plate (ThermoFisher) in E8 medium containing 10 μM Y-27632 (#Y0503, Sigma-Aldrich). The medium was switched to E8 without Y-27632 the following day. Individual colonies were isolated and expanded in E8 medium. To identify targeted clones, colonies were screened by allele-specific PCR using primers SCN2AgcR (5-gagactttggaggggttACT-3) and SCN2AscreenF (5-ctatgttaagagggaagttgg-3). Following multiple rounds of subcloning a single colony was found to contain targeted *SCN2A* gene correction. Expansion and subsequent culture of both corrected lines were performed as per mutant iPSC lines. These cells were referred to as R1882Q_Corrected and R853Q_Corrected iPS cells.

### Lentivirus production

Lentivirus carrying NGN2 and the reverse tetracycline transactivator (rtTA) gene was prepared by first plating 4×10^6^ HEK cells in a T-75 and grown in 5% Foetal Bovine Serum (#FFBS-500, Scientifix) in DMEM/F12 (#10565018, Gibco). The following day cells were transfected using Lipofectamine 2000 (#11668027, ThermoFisher) with plasmid DNA from either FUW-TetO-Ngn2-P2A-puromycin (#52047, Addgene) or FUW-M2rtTA (#20342, Addgene) and pMDL (#12251, Addgene), vSVG (#8454, Addgene), RSV (#12253, Addgene), at a DNA molarity ratio of 4:2:1:1, respectively. Viral supernatant was collected at 24, 48 and 72 hours, filtered through a 0.45μm membrane (#SLHV033RS, Millipore, USA) and concentrated by centrifugation at 85,000x g for 2 hours at 4°C in a Sorvall WX 100 Ultra Ultracentrifuge (ThermoFisher). The supernatant was discarded and viral pellet resuspended in a volume of PBS containing calcium and magnesium (#14090-055, ThermoFisher) that resulted in a 200-fold enrichment. Typically, 0.25-0.5 μl of concentrated virus was used for each neural differentiation, as determined by viral titration.

### NGN2-based neural differentiation

For this study cortical excitatory neurons were generated by the expression of *NGN2* in R1882Q, R853Q and both corresponding corrected iPS cells, as described by (22). Accordingly, at 4 DIV, neurons were lifted to coverslips in a 24-multiwell plate in preparation for quantitative real time PCR (RT-qPCR), immunostaining analysis and single cell electrophysiological recording or onto MEAs (see corresponding section).

### RT-qPCR

Total RNA was isolated using PureLink RNA Mini Kit (#12183018A, ThermoFisher) according to the manufacturer’s instructions. The synthesis of single strand cDNA and quantification of each mRNA was performed using the TaqMan RNA-to Ct 1-step Kit (#4392938, ThermoFisher) a ViiA Real-Time PCR System (Applied Biosystems). Experiments were conducted in multiplexing to obtain relative levels of each transcript normalized for the endogenous controls *GAPDH, HMBS* and *B2M* in every sample. For the RT-qPCR analysis of *MAP2*, given a detected interference with *GAPDH* probe, the endogenous control *GUSB* was used. Each reaction was run in triplicate and contained 20 ng of RNA in a final reaction volume of 10 μL. The specific probes (ThermoFisher) that have been used are as follow: *B2M* (Hs00187842_m1), *GAD1* (Hs01065893_m1), *GAPDH* (Hs99999905_m1), *GUSB* (Hs00939627_m1), *HMBS* (Hs00609297_m1)*, MAP2* (Hs00258900_m1), *SCN1A* (Hs00374696_m1), *SCN2A*(Hs01109871_m1), *SCN3A* (Hs00366902_m1), *SCN8A* (Hs00274075_m1), SCN8A (adult transcript variant) (Hs01578884_m1), and VGLUT *(SLC17A7*,Hs00220404_m1). Data are shown as mean ± SD of n=3 technical replicates obtained from n=3 separate biological experiments (differentiations).

Statistical analysis was performed using GraphPad Prism 8 software. For the two independent sample comparison, the statistical significance was determined using the Holm–Sidak t-test and alpha=5.000%.

### Immunocytochemistry

Cells were fixed in cold 4% paraformaldehyde (Australian Biostain) on ice and permeabilized using 0.2% (v/v) Triton X-100 (Sigma) in PBS (ThermoFisher). Cells were subsequently incubated with 1% foetal bovine serum (#FFBS-500, Scientifix)-PBS^−/−^ blocking solution and stained with primary and secondary antibodies in blocking solution. Immunoreactivity was tested against *SCN2A* (Na_v_1.2) (#AGP-026, Alomone; 1:250), and MAP2 (#ab5622, Abcam; 1:500). For assessing pluripotency of our iPSC lines, immunoreactivity was tested against NANOG (#ab21624, Abcam, 1:100) and TRA-1-60(R) (#ab16288, Abcam, 1:100).

The secondary antibodies were: Alexa Fluor-conjugated 488 anti-Mouse (#A11001, Life Technologies), Alexa Fluor-conjugated 594 anti-Rabbit (#A21207, Life Technologies), Alexa Fluor-conjugated 488 anti-Rabbit (#A11008, Life Technologies) and Alexa Fluor-conjugated 647 anti-Guinea Pig (#CF647, Sigma-Aldrich). Nuclei were visualized using 4,6-diamidino-2-phenylindole, dihydrochloride (DAPI) counterstain (1 mg/mL; #D9452-50MG, Sigma-Aldrich). Samples were mounted onto glass slides using Prolong Gold mounting media (#P36934, ThermoFisher). Images were acquired using a Nikon A1R or Zeiss 780 confocal microscope.

### Whole-cell patch clamping

Neurons were differentiated as described above and grown on 12 mm coverslips (#L2020, Sigma-Aldrich) pre-coated with 100 μg/ml poly-D-lysine (#P7280, Sigma-Aldrich) followed by 15 μg/mL laminin. Neurons were superfused with BrainPhys^™^ Neuronal Medium (Stemcell Technologies, Australia) at 30 ± 0.5°C with a flow rate of 1 mL/min. For phenytoin studies (Figure 4), stock solutions (100 mM in DMSO) of phenytoin (#P1300000, Sigma-Aldrich) were diluted in BrainPhys Neuronal Medium and perfused during patch clamp recordings. All solutions (including vehicle) were matched for DMSO concentration. Thick-walled borosilicate pipettes (4-6 MΩ; #30-0060, Warner Instruments) were filled with solution containing (in mM): 130 K-gluconate, 10 D-glucose, 6 KCl, 5 EGTA, 5 HEPES, 4 NaCl, 2 MgATP, and 0.3 GTP-tris salt (pH 7.3 with KOH, 290 mOsm). Data were recorded with a Multiclamp 700B amplifier controlled by Clampex 10.6/DigiData 1550B acquisition systems (all from Molecular Devices) at a sampling rate of 10 kHz. Resting membrane potentials and spontaneous action potential (AP) firing were recorded using I=0 gap-free protocol for > 30 seconds. Passive and active membrane properties were measured from voltage responses generated using a current clamp protocol in which the cell was held at −70 mV and injected with current from −10 to 135 or 235 pA at 5 pA intervals for 1 sec. Analyses were performed using Clampfit 10.6 (Molecular Devices, USA). GraphPad Prism 8 (GraphPad, USA) was used for graphing and statistical analyses. Data contain recordings from at least three independent differentiations with the patient and corresponding isogenic control cell lines generated and recorded in parallel for each differentiation.

### Soma measurement assay

To compare the soma size of neurons derived from R853Q iPSC cell line with its corresponding control, cells were stained using the CellMask Plasma Membrane Stain following manufacturer’s instructions (#C37608, Life Technologies). Images were obtained from three independent experiments and captured using a Nikon A1R. ImageJ software was used to measure cell soma and GraphPad Prism 8 software for the statistical analysis. For the two independent sample comparison, the statistical significance was determined using the with Holm–Sidak t-tests and alpha=5.000%.

### Multiple Electrode Array (MEA) Analysis

A multiple electrode array system was used to explore the R1882Q mutation at the neural network level. *NGN2* neurons were induced and maintained as described above. 1×10^5^/well *NGN2* neurons and 2.5×10^4^/well primary human astrocytes (#1800, ScienCell) were plated to 24-well 128-electrode MEA plates (#24W700/100F-288, Multi Channel Systems, Germany). Plates have been treated overnight with Terg-a-zyme detergent (#Z273287, Sigma-Aldrich), plasma cleaned (Femto Model Plasma Cleaner, Diener Electronic, Germany), and pre-coated with 100 μg/ml poly-D-lysine (#P7280, Sigma-Aldrich) followed by 15 μg/ml laminin coating (#L2020, Sigma-Aldrich). Medium was changed at least 1 day prior to recordings. Plates were recorded for 10 minutes at 37°C with the Multiwell MEA System and Multiwell Screen software (Multi Channel Systems, Germany). Three independent biological replicates, with patient and corrected cell lines running in parallel for each replicate, were recorded. Spike and burst detection features were extracted as described previously (61). All parameters reported were mean values from each well. Mean firing rate and network burst rate values were calculated from all wells while mean jitter and kappa were derived only from bursting wells. Jitter is a measure of how simultaneous a burst is and is the time from the start of the first burst pike and the start of the last burst spike. Kappa is an indication of the synchronization between spikes. It is calculated between pairs of channels as kappa = (P_obs_ – P_exp_)/(1 – P_exp_), where Pexp is the expected probability of two channels having the same state (spiking or not spiking) randomly and Pobs is the observed probability of the two channels having the same state. The expected probability P_exp_ = (n_i0_n_j0_ + n_i1_n_j1_)/N and the observed probability Pobs = (n_ij0_ + n_ij1_)/N, where N is the total number of bins (10 ms bins used), n_i0_ and n_j0_ are the number of bins where there is no spiking in the i^th^ and j^th^ channels respectively, n_i1_ and n_j1_ are the number of bins with spikes in the i^th^ and j^th^ channels respectively, n_ij0_ is the number of bins where both channels have no spikes and n_ij1_ is the number of bins where both channels are spiking. As kappa is a ratio of probabilities, it takes a value between 0 and 1, with values closer to 0 and 1 indicating less or more synchronization respectively.

### RNA sequencing and pathway analysis

To identify genes and pathways that may be dysregulated in DEE *SCN2A* variants, mRNA sequencing was performed. *SCN2A* R1882Q and R853Q patient and control derived neurons were collected at DIV 21 and processed for RNA sequencing using the PureLink RNA Mini Kit (#12183018A, ThermoFisher) according to the manufacturer’s instructions. RNA samples from 3 biological replicates underwent DNase treatment using DNA-free DNA removal Kit (Cat #AM1906, ThermoFisher) and quantified using the NanoDrop 2000. Samples were sent to the Australian Genome Research Facility (AGRF, Melbourne, Australia) for high-depth mRNA poly(A) sequencing (~100 million mapped reads per sample, 150bp paired-end reads, stranded library protocol) using the NovaSeq Illumina Sequencing platform.

Read quality was assessed using the FastQC software version 0.11.9 (www.bioinformatics.babraham.ac.uk/projects/fastqc/). Alignment and quantification of total RNA-sequencing data was performed using Rsubread aligner (version 2.4.3). Paired-end 150bp Illumina reads were aligned to the human genome (Ensembl GRCh38 assembly) at the gene-level with an average of 97% (±0.19 SD) of reads successfully mapped. Genes with greater than one count-per-million mapped reads in at least two samples were retained for further analysis, genes below this threshold were filtered out.

Differential gene expression analysis was performed in the R statistical programming environment (version 4.0.5) using EdgeR (62) (version 3.32.1), which applied adjustment for library size and normalisation with trimmed mean of M values (TMM) where dispersion parameters for each gene are estimated with the Cox-Reid common dispersion method and employed in a negative binomial generalized linear model for each gene. Accounting for gene dispersion ensures that expression differences that are consistent between replicates are more highly weighted than those that are not to ensure differential expression is not driven by outliers. P-values were adjusted for multiple testing using the Benjamini-Hochberg correction with an FDR q-value <0.05 cut off.

To obtain an insight into the likely functional relevance of differentially expressed genes we undertook a gene ontology (GO) enrichment analysis with the GOrilla web application (63). All genes (ranked by FDR q-value) were uploaded to GOrilla for each study and were used to identify gene ontology terms (biological processes, molecular functions, cellular components) potentially relevant to mutation effects. GO term p-values were corrected for multiple testing with FDR adjustment.

Statistical analysis was described individually for each section above.

This study was approved by the Austin Health Ethics committee, ethics number HREC/16/Austin/472.

## Acknowledgements

This study was supported by a National Health and Medical Research Council (NHMRC) program grant (10915693) to St.P., Medical Research Future Fund (MRFF) Accelerated Research Stem Cell grant to St.P. and Sn.M. and MRFF Genomic Health Futures Mission Project Grant to Sn.M., G.B., and St.P., and project funding by RogCon, Inc. We thank patients and their families for the participation in the study. We thank Dr Marius Wernig for providing *NGN2* plasmids and Lucas Teasdale for proofreading the manuscript.

## Author Contributions

M.M., C.M. and B.R. designed and conducted experiments and analyzed data. S.B. performed bioinformatic analysis. C.C. conducted quality control experiments and analyzed data. G.B. performed preliminary studies. J.H., Sv.P., D.A., L.J., and T.M. performed and contributed to experiments. K.D. & A.N. contributed to the study design. Sa.M. recruited and assessed patients. Sn.M. and St.P. designed the study. M.M., C.M., Sn.M. and St.P. wrote the manuscript.

