## Supplementary materials for "Distinctive *in vitro* phenotypes in iPSC-derived neurons from patients with gain- and loss-of-function *SCN2A* developmental and epileptic encephalopathy"

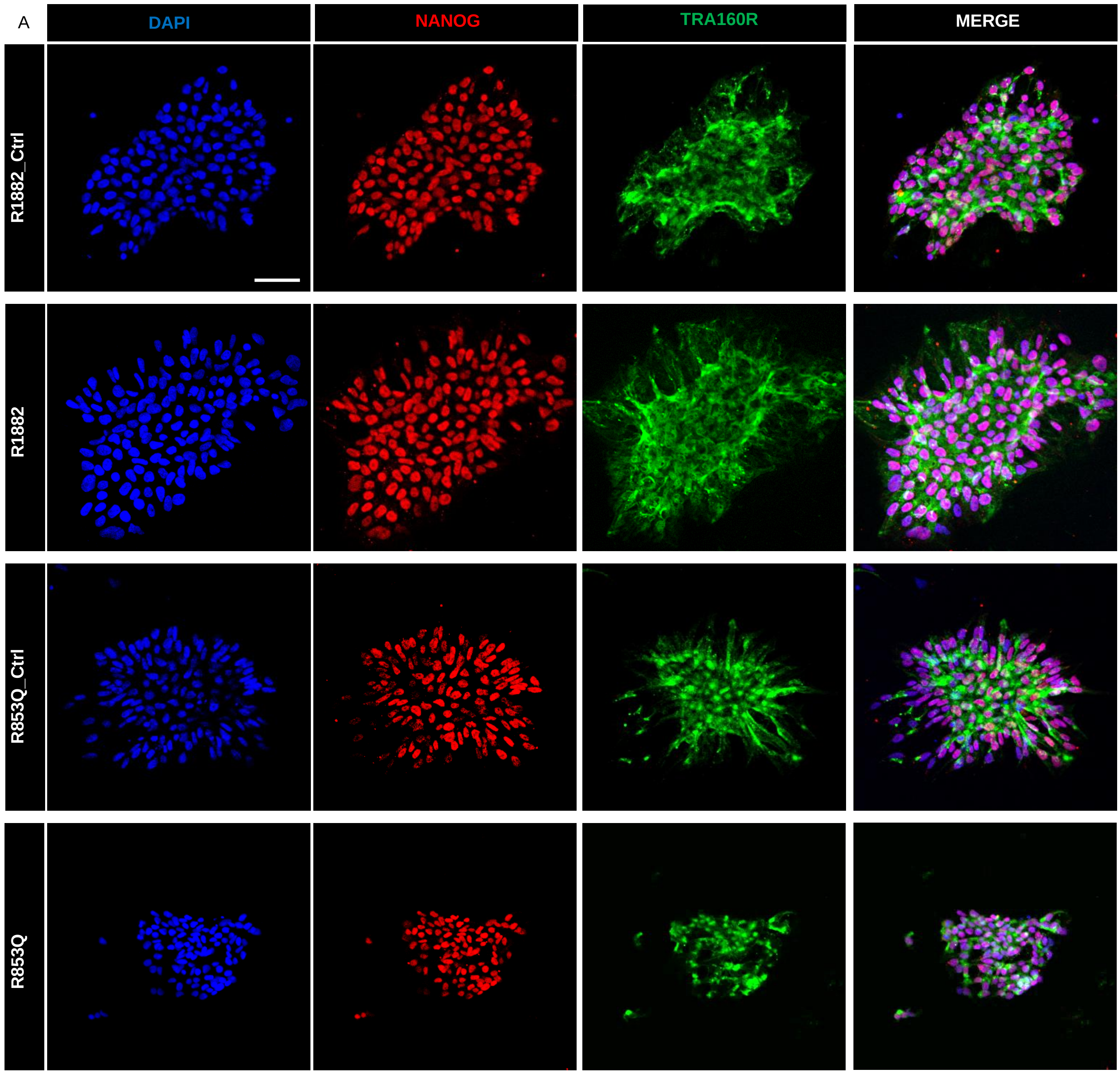


**Figure S1. Characterization of *SCN2A* R1882Q and R853Q variant iPSCs.** (A) Immunoreactivity of patient iPSCs and corresponding isogenic controls to pluripotency markers NANOG and TRA160R. Scale bar, 50 µm.

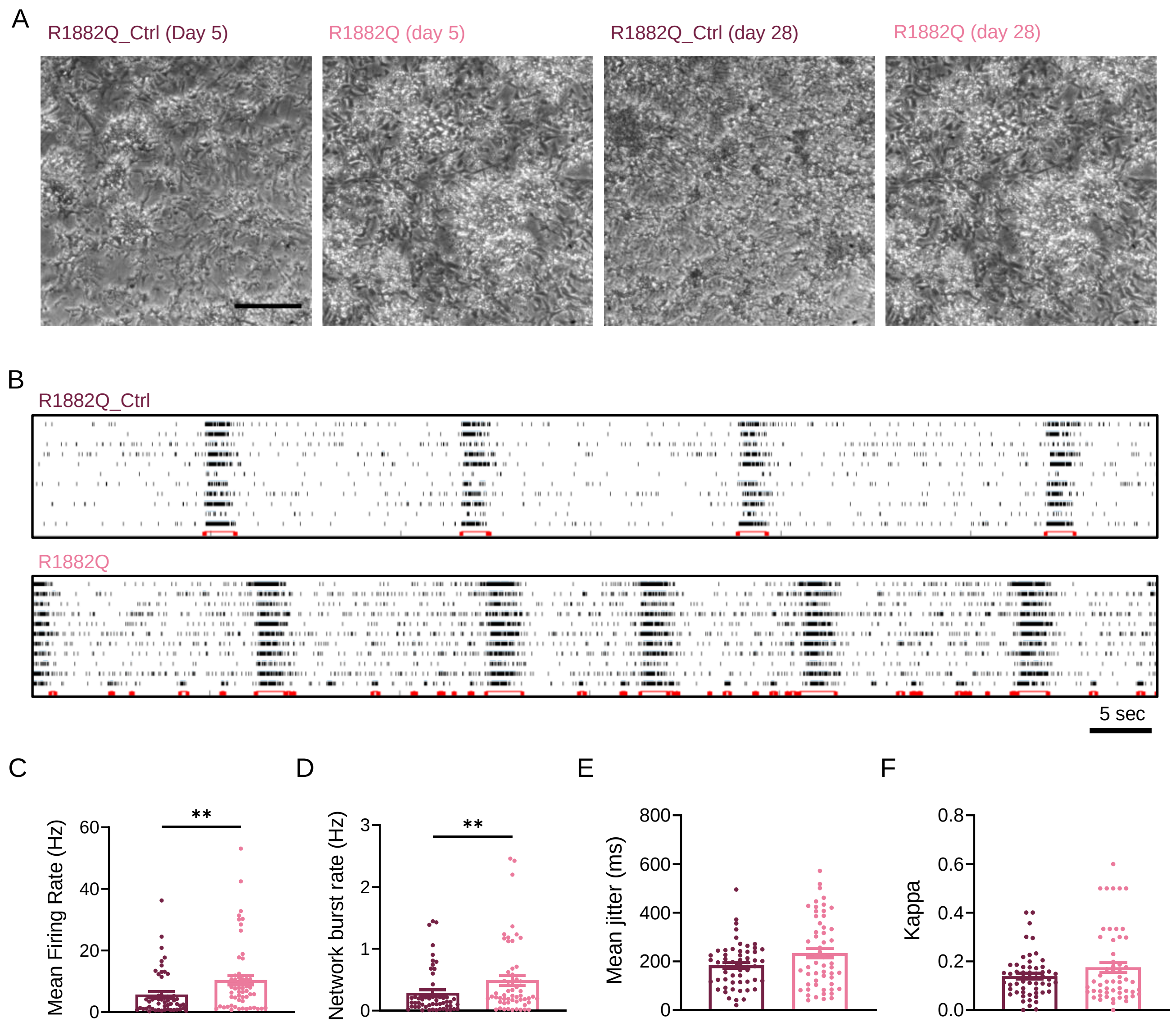


**Figure S2. Multi-electrode array recordings of R1882Q and isogenic control cell lines.** (A) Representative images showing *NGN2* differentiated neurons and astrocyte co-culture at two time points. Scale bar = 200 µm. (B) Representative raster plots from a single well of MEA recordings. Each row represents a single electrode. Black lines and red bars indicate spikes and network bursts, respectively. (C) Mean firing rate ($p$=0.0016). (D) Network burst rate ($p$=0.009). (E) Mean jitter ($p$=0.2025). (F) Kappa ($p$=0.7881). Unpaired Mann-Whitney tests (A, E and F). Mean±SEM.

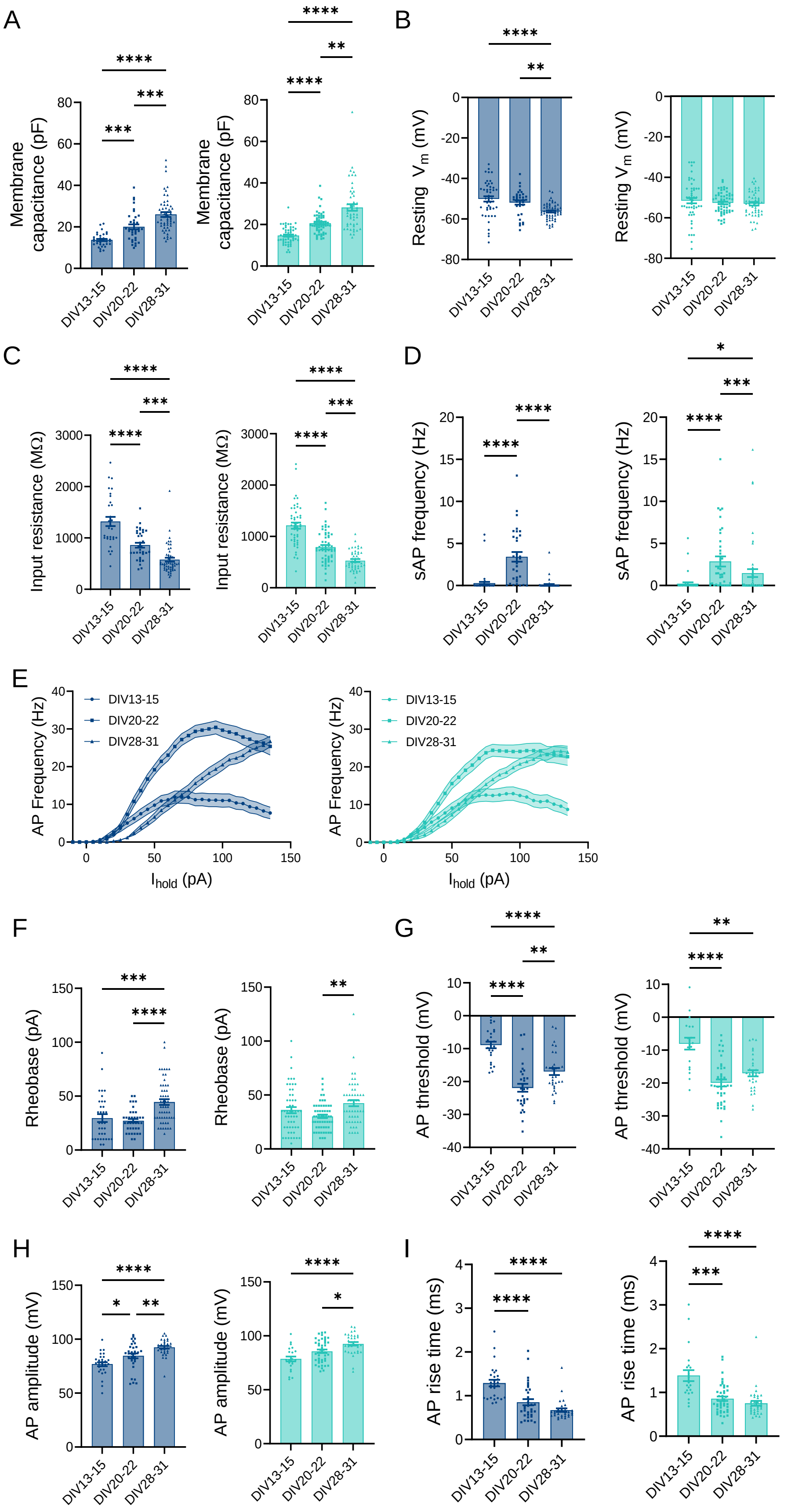


**Figure S3.** ***In vitro*** **development of electrophysiological properties in R853Q (light blue) and corresponding isogenic control (dark blue) iPSC-derived neurons.** (A) Membrane capacitance. (B) Resting membrane potential. (C) Input resistance. (D) Spontaneous AP frequency. One-way ANOVA with Tukey's multiple comparisons test. (E) Mean input-output function of AP firing in response to current injection. (F) Rheobase. (G) AP threshold. (H) AP peak amplitude. (I) 10-90% AP rise time. One-way ANOVA with Tukey's multiple comparisons test. Mean±SEM.

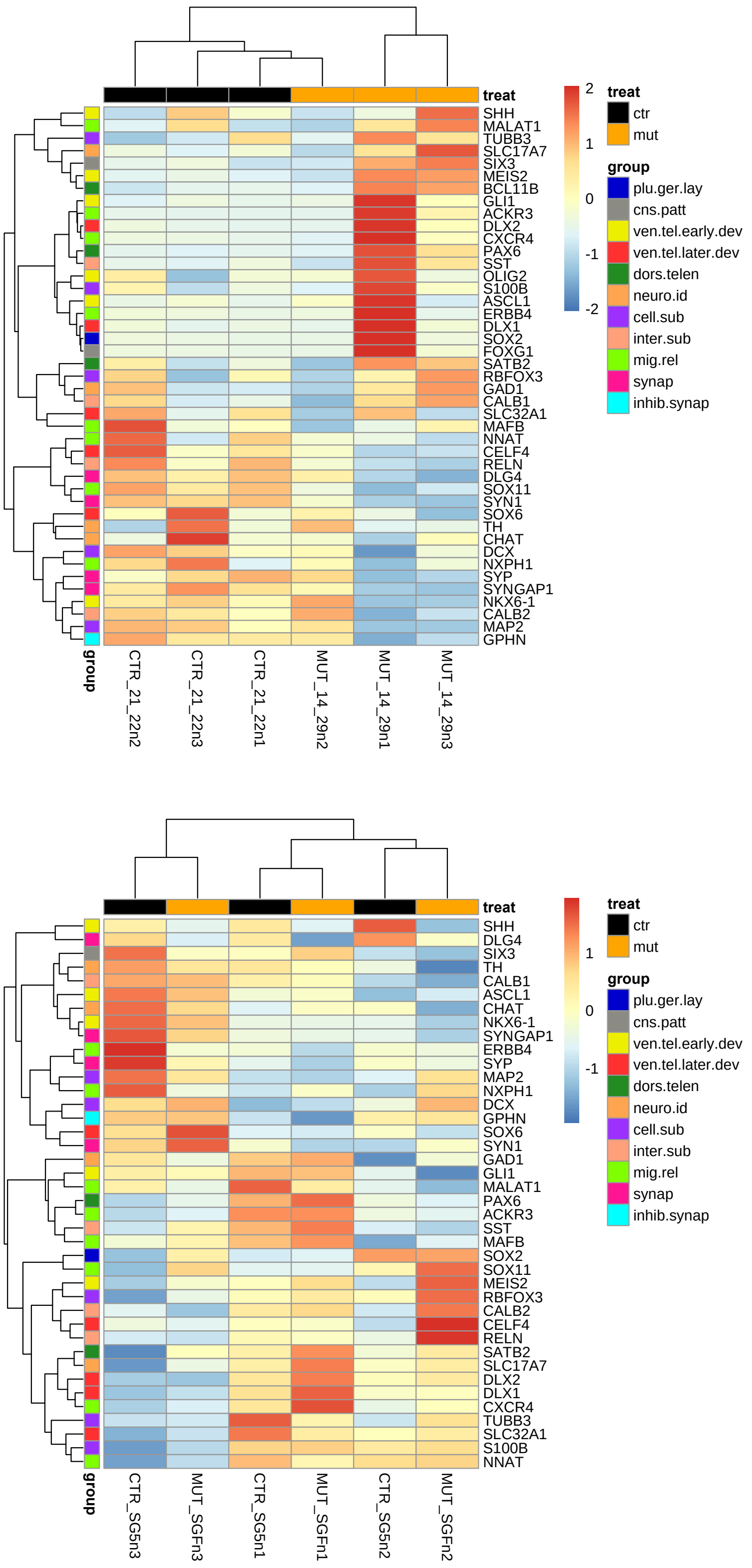


**Figure S4: Gene expression heatmaps and dendrograms for various neural development gene panels in R1882Q (top) and R853Q (bottom) variants based on mRNA-sequencing data.** Gene expressions are scaled (z-score) reads per kilobase of exon per million reads mapped (RPKM) values (main colour key) separated by samples (x-axis) and genes (y-axis). Second and third colour keys represent treatment (control, patient) and gene panels (pluripotency and germ layer (per.ger.lay); CNS patterning (cns.patt); ventral telencephalon (early development) (ven.tel.early.dev); ventral telencephalon (later development) (ven.tel.later.dev); dorsal telencephalon (dors.tele); neurotrasmitter identity (neuro.id); cell subtypes (cell.sub); interneuron subtypes (inter.sub); migration-related genes (mig.rel); synaptogenesis (synap); inhibitory synaptogenesis (inhib.synap). Dendrograms are based on Euclidean distances between genes or samples.

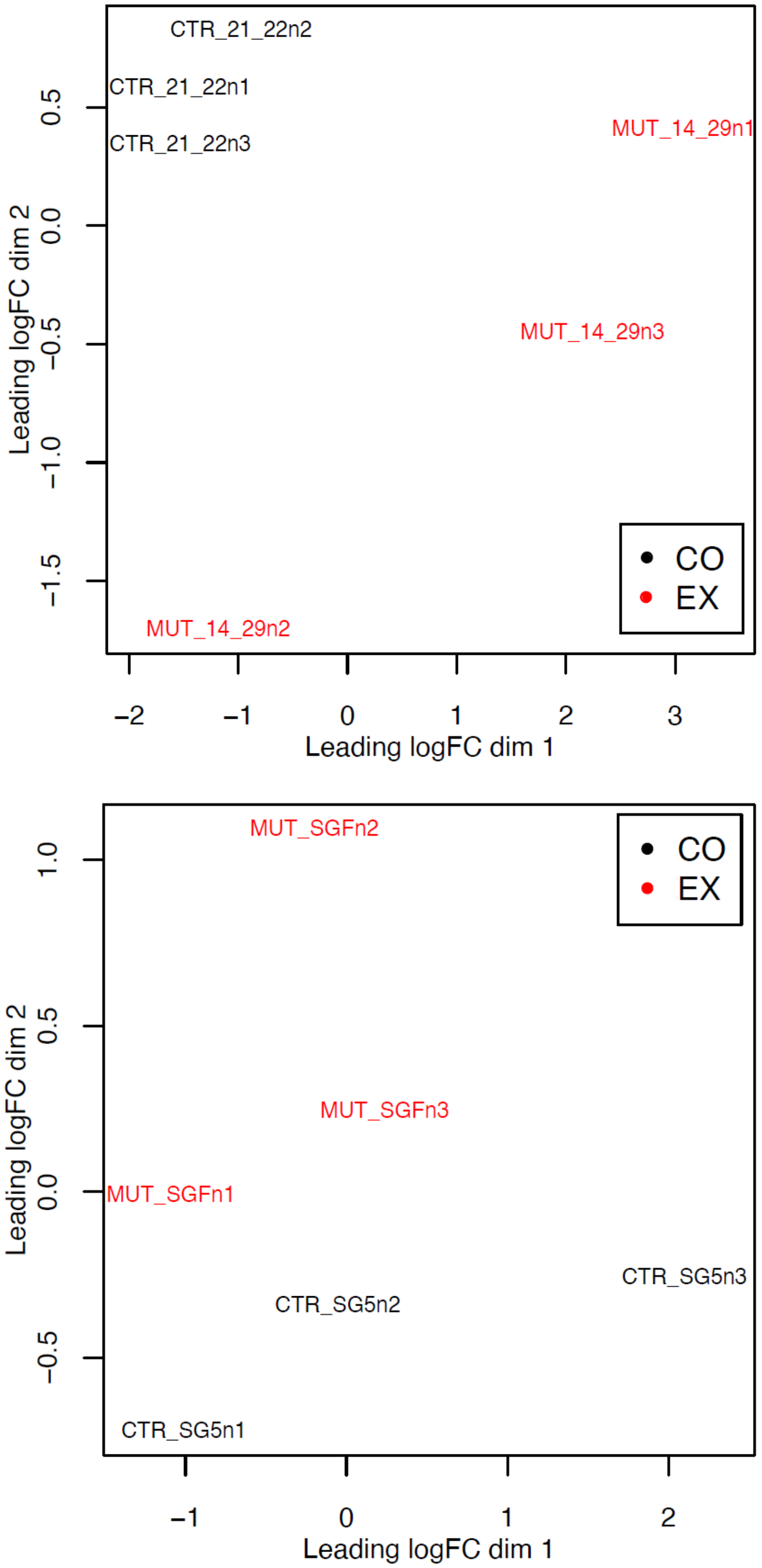


**Figure S5: Multidimensional scaling (MDS) plots showing unsupervised clustering of control and mutant samples in R1882Q (top) and R853Q (bottom) variants based on mRNA-sequencing data.** Distance between each pair of samples is based on root-mean-square of the largest (i.e. 500 DEGs) $log_2FC$ gene expression changes between that pair of samples.

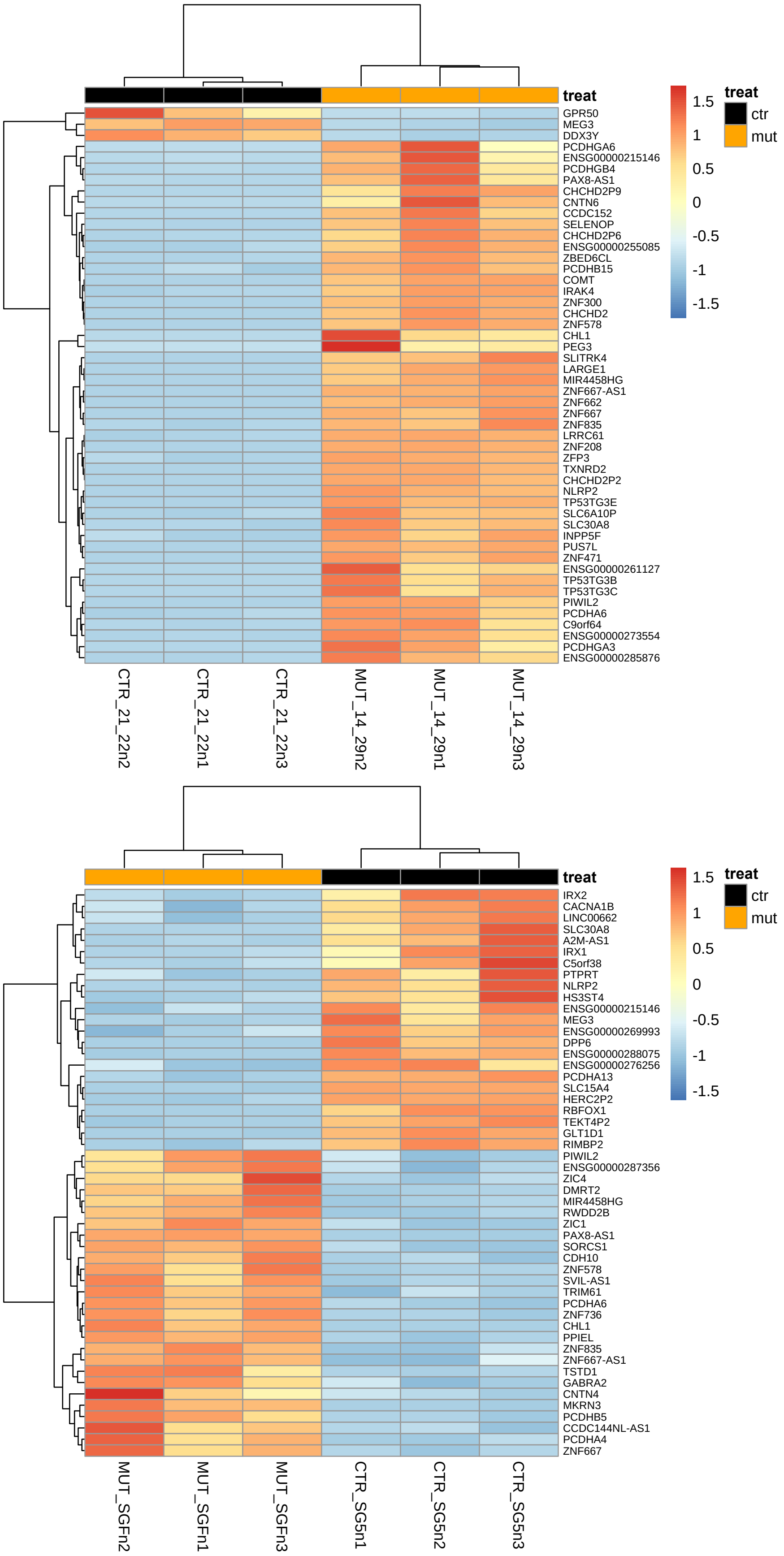


**Figure S6: Gene expression heatmaps and dendrograms for the top 50 significantly differentially expressed genes in R1882Q (top) and R853Q (bottom) variants based on mRNA-sequencing data.** Gene expressions are scaled (z-score) reads per kilobase of exon per million reads mapped (RPKM) values (main colour key) separated by samples (x-axis) and genes (y-axis). Second colour key represents treatment (control, mutant) and dendrograms are based on Euclidean distances between genes or samples.

**Table 1. Summary of baseline electrophysiological properties of all cell lines at each time point.** All statistics were performed using unpaired *t*-tests and are compared to the corresponding isogenic control, *$p<0.05$ and **$p<0.01$. Mean±SEM.

| | *DIV20-22* | | *DIV13-15* | | *DIV20-22* | | *DIV28-31* | |
|---|---|---|---|---|---|---|---|---|
| | R1882Q_Ctrl | R1882Q | R853Q_Ctrl | R853Q | R853Q_Ctrl | R853Q | R853Q_Ctrl | R853Q |
| Passive properties | | | | | | | | |
| Membrane capacitance (pF) | 19.37±0.89 | 17.50±0.68 | 13.73±0.55 | 15.96±0.73* | 20.07±1.21 | 19.44±0.58 | 26.03±1.16 | 28.17±1.56 |
| Resting membrane potential (mV) | -52.25±1.12 | -51.97±1.11 | -51.28±1.52 | -50.78±1.67 | -51.83±1.12 | -52.13±0.71 | -56.31±0.54 | -53.06±0.85** |
| Spontaneous AP frequency (Hz) | 2.10±0.58 | 1.17±0.39 | 0.40±0.24 | 0.36±0.21 | 3.42±0.57 | 2.87±0.58 | 0.12±0.08 | 1.47±0.48** |
| Input resistance (MΩ) | 918.2±56.7 | 815.5±56.8 | 1318±92.8 | 1169±74.0 | 862.1±47.4 | 786.1±47.9 | 576.3±38.6 | 529.1±29.2 |
| AP properties | | | | | | | | |
| Rheobase (pA) | 27.35±1.83 | 32.96±2.41 | 29.56±3.58 | 29.85±3.51 | 27.16±1.77 | 31.05±2.02 | 44.44±2.76 | 42.45±2.84 |
| Threshold (mV) | -17.39±1.14 | -21.24±0.97* | -8.92±0.97 | -8.07±1.78 | -21.91±1.24 | -21.22±1.05 | -16.93±1.08 | -17.01±0.95 |
| Peak amplitude (mV) | 79.96±1.14 | 83.52±1.31* | 77.00±1.87 | 78.51±2.36 | 84.56±2.18 | 86.01±1.85 | 92.42±1.35 | 92.49±1.63 |
| Half-width (ms) | 1.27±0.03 | 1.32±0.04 | 1.50±0.07 | 1.54±0.10 | 1.31±0.07 | 1.31±0.05 | 1.38±0.06 | 1.39±0.04 |
| 10-90% rise time (ms) | 0.92±0.05 | 0.78±0.04* | 1.29±0.07 | 1.39±0.12 | 0.85±0.07 | 0.83±0.06 | 0.67±0.04 | 0.75±0.06 |
| 90-10% decay time (ms) | 1.83±0.08 | 1.88±0.06 | 2.22±0.14 | 2.27±0.17 | 2.00±0.10 | 2.08±0.11 | 2.13±0.12 | 2.20±0.11 |
| Maximum frequency (Hz) | 38.71±1.25 | 42.89±1.29* | 26.29±1.16 | 25.97±1.52 | 42.22±1.21 | 41.56±1.34 | 27.56±1.34 | 27.37±1.26 |

**Table 2. Summary data for effect of phenytoin on AP firing in R1882Q and R853Q cell lines.** All statistics were performed using one-way ANOVA with Tukey's multiple comparisons test, $^{**/\#\#}p<0.01$ and $^{****/\#\#\#\#}p<0.0001$. *, compare to R1882Q isogenic control (Vehicle); $^{\#}$, compare to R1882Q patient (Vehicle). Mean±SEM.

| *Cell line/Treatment* | *Max. AP Frequency (Hz)* | *Area Under Curve (A.U.)* |
|---|---|---|
| R1882Q | | |
| R1882Q isogenic control (Vehicle) | 33.3±1.4 | 3448±236 |
| R1882Q patient (Vehicle) | 40.5±1.4** | 5067±271**** |
| R1882Q patient (30 µM) | 29.9±3.4$^{\#\#}$ | 3187±441$^{\#\#}$ |
| R1882Q patient (70 µM) | 21.4±2.7$^{**/\#\#\#\#}$ | 2227±322$^{\#\#\#\#}$ |
| R1882Q patient (100 µM) | 17.9±1.7$^{****/\#\#\#\#}$ | 1748±210$^{**/\#\#\#\#}$ |
| R853Q | | |
| R853Q isogenic control (Vehicle) | 31.9±2.2 | 2961±431 |
| R853Q patient (Vehicle) | 25.8±3.2 | 2735±485 |
| R853Q patient (30 µM) | 25.0±2.2 | 2172±263 |
| R853Q patient (70 µM) | 18.6±2.9** | 1788±338 |
| R853Q patient (100 µM) | 12.8±2.2$^{****/\#\#}$ | 948±159$^{**/\#\#}$ |

**Table 3. Description of DEGs in R1882Q and R853Q neurons, and in overlapping comparison**. Summary of gene role and protein function, and relevant bibliography to association with neurological disorders for top three DEGs in R1882Q (*LARGE1*, *ZNF300*, *CHCHD2*) and R853Q (*DPP6*, *GLT1D1*, *MKRN3*) neurons and overlapping DEGs (*CHL1*, *GPR50*, *NLRP2*).

| | GENE | SUMMARY | ASSOCIATION WITH NEUROLOGICAL DISORDERS |
|---|---|---|---|
| R1882Q | *LARGE1* | This gene encodes a member of the N-acetylglucosaminyltransferase gene family, the glycotransferase, that adds the final xylose and glucuronic acid to alpha-dystroglycan and thereby allows alpha-dystroglycan to bind ligands including laminin 211 and neurexin. | Mutations in this gene cause several forms of congenital muscular dystrophy characterized by abnormal glycosylation of alpha-dystroglycan, severe structural brain and eye malformations, and cognitive disability (1,2). |
| | *ZNF300* | This gene encodes a C2H2-type zinc finger DNA binding protein and likely transcriptional regulator (3). The function of this protein is not yet known. | Unknown |
| | *CHCHD2* | This gene protein encodes a eukaryotic CX(9)C protein. In response to stress, the protein translocates from the mitochondrial intermembrane space to the nucleus where it binds to a highly conserved 13 nucleotide oxygen responsive element in the promoter of cytochrome oxidase 4I2, a subunit of the terminal enzyme of the electron transport chain. In addition, it has been shown that this protein is a negative regulator of mitochondria-mediated apoptosis. | Downregulation of *CDCHD2* has been associated with Parkinson's Disease, Alzheimer's Disease and Frontotemporal Dementia (4). |
| R853Q | *DPP6* | This gene encodes a single-pass type II membrane protein that is a member of the peptidase S9B family of serine proteases. This protein binds specific voltage-gated potassium channels and alters their expression and biophysical properties. | *DPP6* loss induces behavioral impairment and impacts hippocampal synaptic development (5) and may cause dysfunction of neuronal excitability in neurodegenerative dementia (6) and Schizophrenia (7). |
| | *GLT1D1* | This gene product is predicted to enable glycosyltransferase activity. It is located in cytosol. | Unknown |
| | *MKRN3* | This gene product contains a RING (C3HC4) zinc finger motif and several C3H zinc finger motifs. This gene is intronless and imprinted, with expression only from the paternal allele. | Disruption of the imprinting at this locus may contribute to hypothalamic disfunction in Prader-Willi syndrome (8). |
| OVERLAP | *CHL1* | This gene encodes a member of the L1 family of neural cell adhesion molecules. It is highly expressed in the central and peripheral nervous system playing an important role in the building and functioning of the brain. | Deletions of this gene have been described in patients with neurodevelopmental delay and seizures. Less is known about the potential effect of *CHL1* overexpression, and microduplications of *CHL1* have been rarely identified. Notably, two patients with a *CHL1* copy number variation (CNV) were reported and both presented some overlapping phenotypes including intellectual disability (9,10). |
| | *GPR50* | This gene product belongs to the G-protein coupled receptor 1 family. | Polymorphic variants of this gene have been associated with bipolar affective disorder in women (11) and ASD (12). |
| | *NLRP2* | This gene product is a member of the nucleotide-binding and leucine-rich repeat receptor (NLR) family which is involved in the regulation of immune responses. | Dis-regulation of *NLRP2* has been observed in familial early-onset Alzheimer's (13), late-onset Alzheimer's (14,15), and in bipolar disorder (16,17). |
